# Unlocking DOE potential selecting the most appropriate design for rAAV optimization

**DOI:** 10.1101/2024.05.06.592659

**Authors:** Konstantina Tzimou, David Catalán-Tatjer, Lars K. Nielsen, Jesús Lavado-García

## Abstract

The production of recombinant adeno-associated virus (rAAV) for gene therapy via triple transfection is a highly intricate process involving many cellular interactions. Each of the different elements encoded in the three required plasmids—pHelper, pRepCap, and pGOI— play a distinct role and affect different cellular pathways when producing rAAVs. The expression balance of these different elements emphasizes the critical need to fine-tune the concentration of all three plasmids and transfection reagents effectively.

The use of design of experiments (DOE) to find optimal plasmid and transfection reagent ratios is a powerful method to streamline the process. However, the choice of the DOE method and the design construction is crucial to avoid misleading results. In this work, we examined and compared four distinct DOE approaches: a rotatable central composite design (RCCD), a Box-Behnken design (BBD), a face-centered central composite design (FCCD), and a mixture design (MD). We compared the ability of the different models to predict optimal ratios, interactions among the three plasmids and transfection reagent, and the essentiality of studying the variability caused by uncontrolled random effects using blocking.

Our findings revealed that MD, when coupled with FCCD, outperformed all other tested models. This outcome underscores the importance of selecting a model that can effectively account for the biological context, ultimately yielding superior results in optimizing rAAV production.

## 1. Introduction

Gene therapy is a revolutionary medical technology that involves the *in vivo* introduction, alteration, or deletion of genetic material to treat or prevent diseases. It aims to correct faulty genes, supplement missing or defective ones, or modulate gene expression to restore normal cellular function. Gene therapy has shown great potential addressing a wide range of genetic disorders, such as cancer, blood disorders, muscular dystrophies, or cardiovascular diseases^1^. Its potential lies in providing a treatment of hitherto untreatable diseases or an alternative where existing treatments are accompanied by severe side effects that significantly compromise the patient’s quality of life. Although only 10 gene therapy products have been approved by FDA, there is a high number of gene therapies under clinical trials, paving the way for future personalized medicine^2^.

Several platforms are available for delivering therapeutic genes to the patient, including viral vectors, liposomes, inorganic nanoparticles, and cationic polymers. Recombinant adeno-associated viruses (rAAVs) have gained high attention in recent years as *in vivo* gene delivery vectors, due to their favorable safety profile, high efficiency in gene delivery and broad tropism for specific tissues^3^. The predominant method to produce rAAVs to date involves triple transfection of mammalian cells. Meeting the high demand for rAAVs is impeded by the scalability of this production process and the challenges associated with large-scale transient gene expression (TGE)^4,5^. Optimisation is crucial but can be challenging due to the involvement of multiple parameters, including cell line, cell culture conditions such as culture media and cell density, and transfection conditions such as total plasmid DNA amount, the plasmid ratio employed in the transfection process and the total amount of transfection reagent ^6–10^.

The use of design of experiments (DOE) to study, characterize, and optimize a process is well-established in industry and widespread into multiple and different fields such as food manufacturing, agriculture, material design, even e-commerce and Big Data analysis, among others. This systematic and efficient approach allows researchers to study the effects and interactions of multiple variables simultaneously, minimizing experimental runs and providing statistically robust models for prediction of optimal solutions^11–14^. However, applying DOE to tackle biological problems can be challenging, due to the high variability of biological systems and the complicated nature of interactions of biological components ^2,15^. In the past, optimization of rAAV production was mainly based on one factor at a time (OFAT) methods, which are easier to approach but more time-consuming^16–19^. Recently, DOE approaches started gaining attention as a tool for rAAV production optimization. By systematically varying the different parameters, different DOE approaches lead to an efficient exploration of different experimental spaces^11^. Traditionally, response surface methodologies (RSM) have been employed for modeling and optimizing responses in biological systems. Central composite design (CCD) and Box-Behnken design (BBD) are the most used approaches to optimize biological processes^20–26^. Regarding rAAV optimization, most of these studies also rely on RSM and the factors used in the designs are sometimes not independent, making it challenging to analyze the generated models^27–29^. These models employ second-order polynomials to identify optimal combinations of the studied variables^30^. While a CCD combines factorial points at the extremes of the studied limits of the variables, BBD tests combinations of factors at the midpoints of the limits of the experimental space and lack points at the edges, ensuring that all design points remain within the safe operating zone, avoiding extreme factor settings simultaneously^31^. One of the advantages of using these RSM methods is that they can present rotatability, meaning that all tested points in the experimental space have equal distance from the design center, ensuring a consistent variance prediction in all runs and enhancing the design’s efficiency. A CCD model can always be made rotatable (RCCD), adjusting the limits of some of the runs while a BBD is only fully rotatable for specific designs^31^. Moreover, depending on the biological response, we might need to test both the edges and midpoints of the experimental space. In this case, a CCD can also be made to test both adjusting its axial points to the center, in a face-centered CCD (FCCD). This configuration lacks rotatability and, thus, does not allow for assessment of potential curvature within the system^31^. These designs can be generated and executed in multiple orthogonal blocks, allowing for the independent estimation of model terms and uncontrolled random effects. This approach minimizes variation in regression coefficients, contributing to a more robust and accurate analysis of the experimental factors^32,33^.

Apart from RSM designs, mixture design (MD) is a specialized experimental approach used to study the effects and interactions of different factors when varying their proportions in a mixture, always adding up to a fixed quantity. This dependency on a constant total is a key distinguishing feature from MD compared to RSM, where each factor varies independently. MDs are particularly valuable in fields like chemistry, pharmaceuticals, and food science, where products often consist of a blend of various ingredients, and understanding the optimal ratio combination is crucial^34,35^. Park et al., showed that a MD approach can be used to study rAAV production and plasmid ratios in the triple transfection protocol^27^.

However, these studies differ in the parameters selected as factors and do not compare the outcome and potential of each methodology. In this study, we conducted a comparative analysis of four distinct DOE methodologies: a complete fractional RCCD, FCCD, BBD and MD with the aim to evaluate their differences when studying and optimizing rAAV production. We intend not only to find optimal parameters for the rAAV triple transfection method with each model, but also to analyze advantages and disadvantages for the use of each type of model, facilitating the identification of the most suitable DOE approach tailored to individual cases.

## 2. Materials and Methods

### 2.1. Cell culture

HEK293SF-3F6 cells from the National Research Council of Canada (NRCC) were cultivated in disposable polycarbonate 125-mL baffled shake flasks equipped with vented caps (Corning® Life Sciences, USA) in 20 mL of HyCell TransFx-H (Cytiva Life Sciences, USA), supplemented with 4 mM GlutaMAX™ (Gibco, Life Technologies, ThermoFisher, USA) and 0.1% Pluronic™ F-68 Non-ionic Surfactant (Gibco, Life Technologies). Cultures were maintained at 37°C and 5% CO_2_ in a Hera Cell 150 incubator (ThermoFisher, USA) with agitation of 130 rpm provided by Celltron (Infors HT, Switzerland) to ensure optimal growth conditions.

Samples were collected for cell count and viability assessment at 24-, 48- and 72-hours post-transfection (hpt) using NucleoCounter®-250 (Chemometec, Denmark) according to manufacturer’s instructions.

### 2.2. Triple transient transfection in HEK293SF-3F6 cells

HEK293SF-3F6 cells were subjected to triple transfection using the following plasmids: PF1451 (pGOI), pXR2 (pRepCap) and pXX6-80 (pHelper). PF1451 (Plasmid Factory, Germany) carries the gene-of-interest (GOI) GFP flanked by inverted terminal repeats (ITR), pXR2 (National Gene Vector Biorepository, USA) contains the Rep and Cap genes from AAV2 and pXX6-80 (National Gene Vector Biorepository, USA) provides the helper genes E2 and E4. HEK293SF-3F6 were seeded at 1 × 10^6^ cells/mL the day prior to transfection to ensure that the cultures were at 2 × 10^6^ cells/mL when the transfection was performed. The transfection mix corresponded to 5% of the total working volume. Briefly, DNA was added to the corresponding volume of culture media, vortexed for 10s, followed by the addition of the necessary amount of the transfection reagent FectoVIR®-AAV (Polyplus, France). The DNA and FectoVIR®-AAV (FV) mix was vortexed three times 3s and incubated in accordance with the manufacturer’s instructions. The ratios between each plasmid and between the DNA and the transfection reagent were determined by the corresponding experimental design.

### 2.3. Design of experiments (DOEs) and models used

In this study, we analyzed the effect of viability, measured in percentage of live cells, on the production of rAAV and we maximized the volumetric productivity (Vp), expressed in vg × mL^-1^ × day^-1^. In order to fit the models, the response was transformed to Log(Vp) as presented in equation 1, using 4 different designs: a rotatable central composite design (RCCD), a Box-Behnken design (BBD), a face-centered central composite design (FCCD) and mixture design (MD).

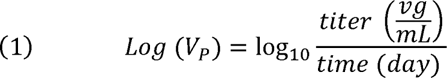

The first three designs (RCCD, BBD and FCCD) treat all 4 factors as independent variables (pHelper, pGOI, pRepCap and FV) to predict optimal plasmid and DNA:FV ratios. MD only includes pHelper, pGOI and pRepCap, presenting the mathematical constraint of the sum of all plasmids being constant, to study plasmid ratios. To study DNA:FV ratio, fixing the optimal plasmid ratio from MD, a two-factor FCCD and RCCD were fit. Considering the nature of the triple transfection system, all designs were constrained to have a minimum quantity of each plasmid and FV to ensure successful production of rAAV.

The experimental design and statistical analysis were conducted using JMP 16 Pro (SAS Institute Inc., USA) for all designs.

#### Response surface methodologies (RSM)

All RSM models (RCCD, BBD and FCCD) fit the experimental space to a second-order polynomial equation (**Equation 2)** where *y* is the response, either Log (Vp) or viability (%), β_0_ is the offset term, β_i_ is the linear coefficient, β_ii_ is the quadratic coefficient, β_ij_ is the interaction coefficient with x_i_ and x_j_ as the independent variable and L as the noise observed in the response.

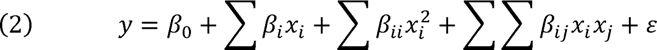

In terms of rotatability, the difference between the three designs lies in the variance of the predicted response at points of interest. If the variance is the same for all points that are at the same distance from the center, the design will be rotatable. To ensure rotatability, the distance from the axial points to the center of the experimental space, α, must be equal to the fourth root of the number of points in the factorial portion of the design (n_F_) to achieve rotatability (**Equation 3).**

For instance, in the 4-factor RCCD presented in this work, n_F_ equals 16 meaning that to have rotatability, α equals 2 while the FCCD has, by definition, α equals 1 and, therefore, it is not rotatable.

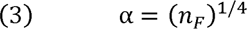

Regarding BBD, all experimental runs lie on a sphere of radius √2 from the center of the experimental space, making BBD rotatable for our 4-factor design.

All RSM designs screened the 4 factors at three levels: a low level coded as −1, an intermediate level coded as 0, and a high level coded as +1 (**Figure 1**).

**Figure 1:**
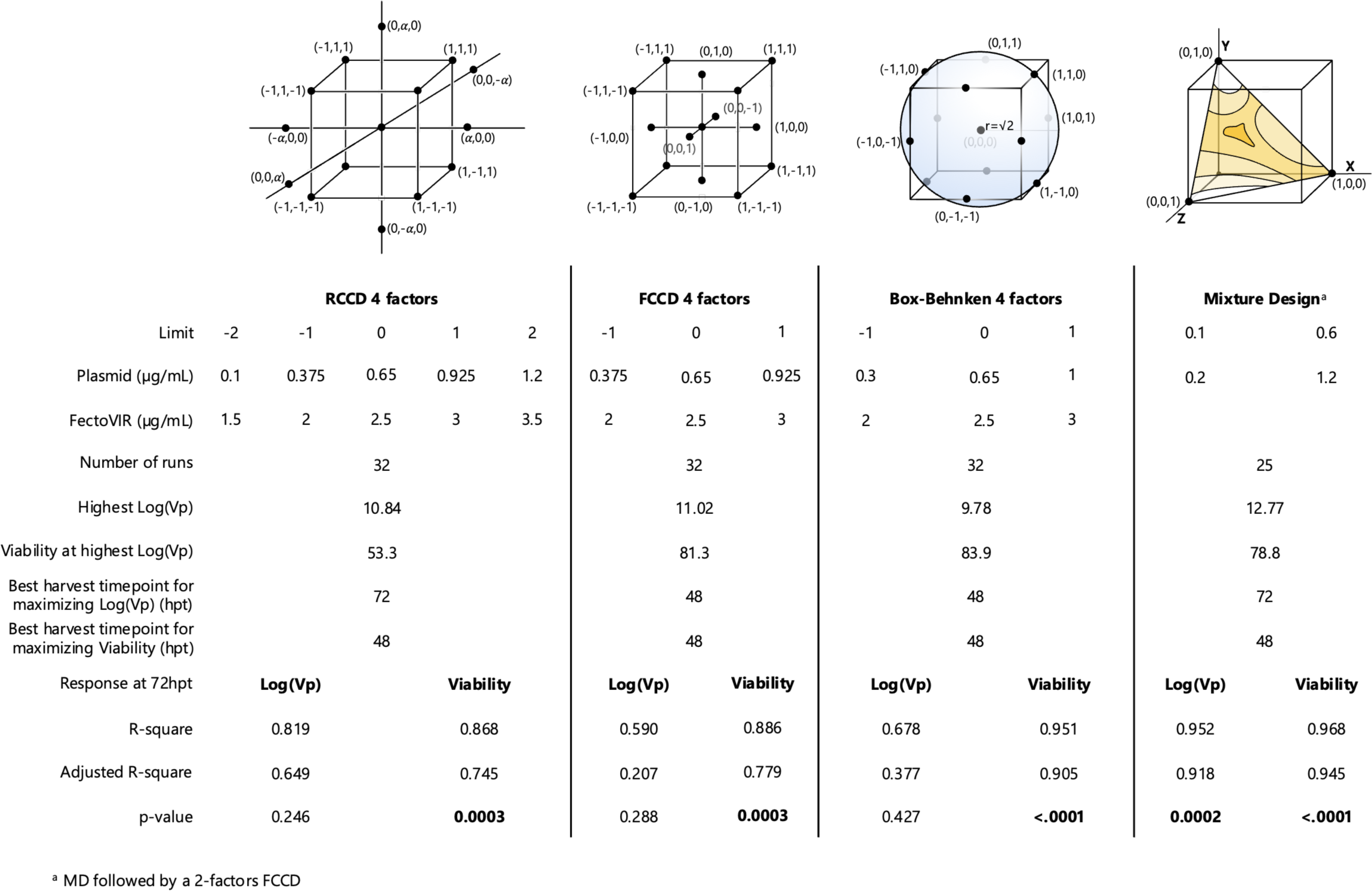
Models comparison. Models comparison between RCCD, FCCD, Box-Behnken design (BBD) and mixture design (MD) with the selected limits, characteristics and statistical parameters for both Log(Vp) and viability responses for each model.

Due to the high complexity of some models, we used the desirability function (**Equation 4**) to identify the optimal values^36^. The desirability function, where *T* is the target, *L* is the lower limit, *y* the response to be maximized and *r* is a number between 0 and 1 to show the relevancy of the response. To analyze more than one response simultaneously, such as viability and Log (V_P_) in our case, we merged both desirabilities in **Equation 5** where *k* is the response.

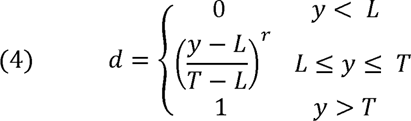

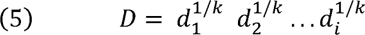

Finally, the contribution of the quantification method to the observed variability was studied as both fixed and random effect for RCCD, BBD and FCCD as they required more than 24 runs and quantification by qPCR was limited to batches of 24 samples simultaneously. To do this, all RSM designs were split into three blocks of 10 runs each with a total number of six central points (2 per block) to ensure the robustness of the design. To analyze the blocks as fixed effect, the significance was calculated by the analysis of variance (ANOVA) test. When the blocks were treated as random effects, the significance of each model was determined by the restricted maximum likelihood method (REML)^37^.

#### Mixture design

The MD was defined as a mixture of the three required plasmids for the successful production of rAAV with a minimum of 10% and a maximum of 60% of each plasmid in the final mixture.

Compared to a cube (RCCD and FCCD) or a sphere (BBD) for the experimental space, in the case of MD we selected an optimal design to place the runs in the experimental space so they minimize the variance of the estimators. This is achieved by minimizing the covariance matrix. This D-optimal criterion is one of the most popular designs and it is available in JMP, Minitab, Design-Expert and other software packages.

All factors were fitted to the quadratic formula presented below **(Equation 6)** where β corresponds to the expected response to the pure blend x_i_ = 1 and x_j_ = 0 when j ≠ i.

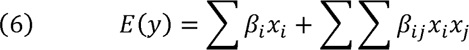

No blocking was required as the number of runs was only 12 and, consequently, the significance of the model was determined by an ANOVA test.

### 2.4. qPCR-based rAAV quantification

Cell culture was harvested at 48 and 72 hpt and diluted 1:1 with lysis buffer (TrisHCl 400 mM, Triton X-100 1% and MgCl_2_ 20 mM adjusted to pH 7.5). After 1h incubation at 37°C and agitation at 130 rpm, the lysate was centrifuged for 20 min at 4000 x g and 4°C. The supernatants were stored at -70°C for long-term storage.

For quantification, samples were thawed in a controlled manner at room temperature. Then, 5 µL of sample were mixed with 2 µL of 1 U/µL DNase I (ThermoFisher Scientific, USA) and 13 µL of 10X DNase I reaction buffer with MgCl_2_ (ThermoFisher Scientific). The mix was incubated for 16 h at 37°C. To inhibit the activity of DNase I, 4 µL of 50 mM EDTA (ThermoFisher Scientific) were added to the mix followed by a 30 min incubation at 70°C. Lastly, 5 µL of Proteinase K (ThermoFisher Scientific) was added to the sample and incubated for 2 h at 55°C. The Proteinase K was inactivated for 15 min at 95°C. 2 µL of the freshly treated sample were mixed with 0.5 µL of 10 µM forward primer (5’-ACGTCAATGGGTGGAGTATTT-3’) and reverse primer (5’-AGGTCATGTACTGGGCATAAT-3’) binding to the GFP sequence with 5 µL of Brilliant III Ultra-Fast SYBR Green QPCR Master Mix (Agilent Technologies, USA) and 0.15 µL of 2 µM reference dye.

Amplification was executed in a QuantStudio 5 Real-Time PCR System (ThermoFisher Scientific) with the following conditions: 50°C for 2 min, 95°C for 10 min; 40×: 95°C for 15 s, 60°C for 1 min, 95°C for 15 s, 60°C for 1 min and 95°C for 1s. Plates run in QuantStudio 5 Real-Time PCR System were analysed via ThermoCloud (ThermoFisher Scientific).

In order to ensure the reliability of the generated data, two controls were added: saturate lysate control (SLC) and transfected lysate control (TLC). The SLC was prepared from a non-transfected cell culture which underwent the lysis protocol in parallel with the transfected samples. Then 5 µL of cell lysate were mixed with 40 ng of pGOI plasmid, 9 µL of 10X DNase I reaction buffer with MgCl_2_ and 2 µL of 1 U/µL DNase I. SLC control aimed to verify that the DNase step was successful. For the generation of the TLC, a single plasmid transfection was performed with pGOI following the triple transfection, lysis and the quantification protocol. The TLC aimed to guarantee that the effect of DNase I remains independent of whether the plasmid has been transfected or spiked afterwards.

## 3. Results and Discussion

### 3.1. Blocking is required when uncontrolled factors are introduced

To study whether the strategy of dividing the runs into three orthogonal blocks was successful, we analyzed the corresponding half-normal plots of each model with each block treated as a fixed factor (**Fig 2B-2E, 3B-3E, 4B-4E**). In these plots, the y-axis represents the normal estimate (orthog t), while the x-axis presents the normal quantile for each RSM (RCCD, FCCD, BBD), each response (Log(Vp), and viability), and each time point (48 and 72 hpt). The blue line intersects the origin with a slope equal to Lenth’s estimate of σ, and any significant effect will not conform to this line. The primary objective was to identify significant effects in each scenario, with a particular focus on whether the blocking strategy was necessary for our system.

**Figure 2:**
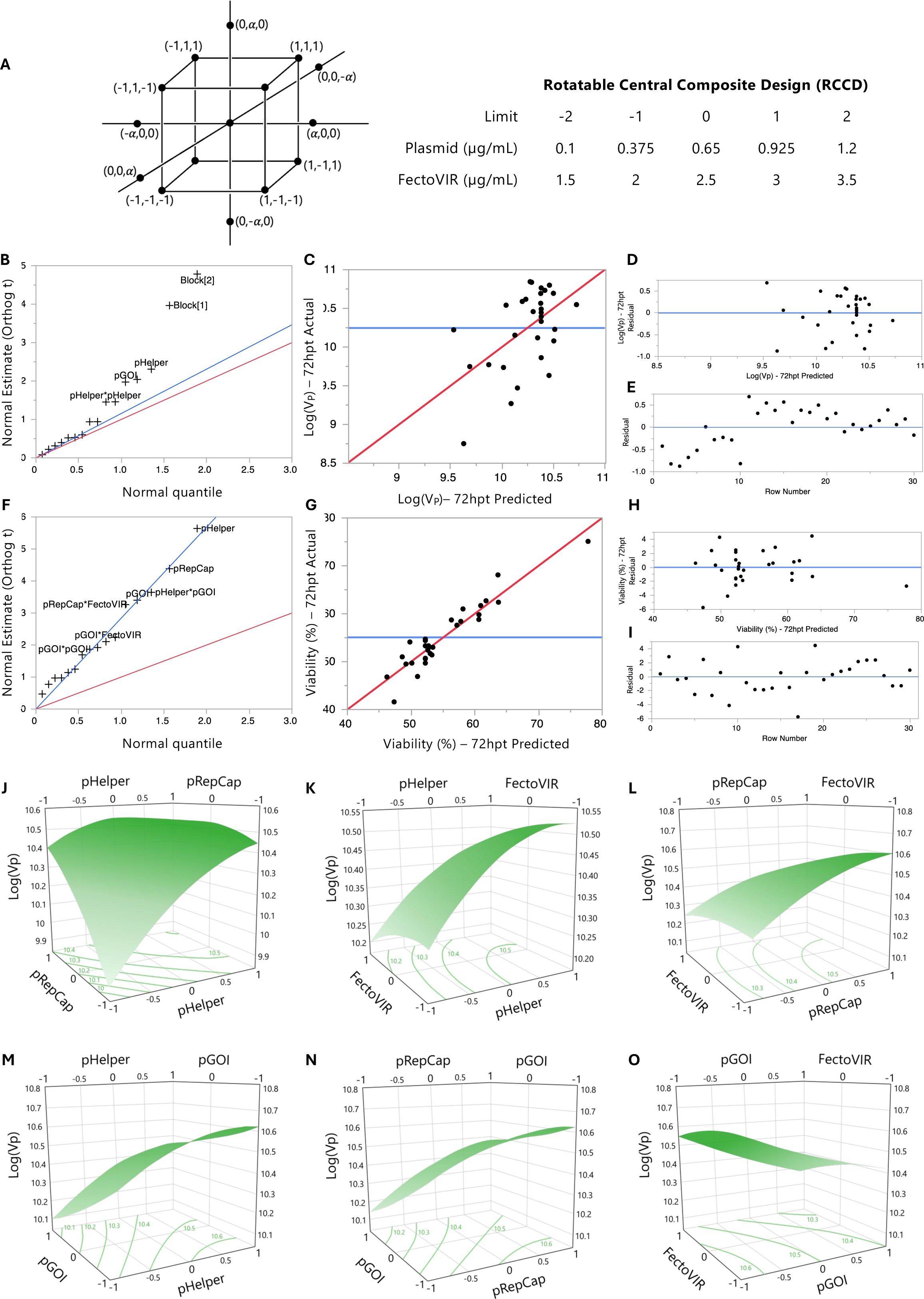
RCCD 72 hpt. (A) Graphic representation of experimental space of a 4-factors RCCD (left) and limits used for the model (right). (B) Normal estimate (Orthog t) against normal quantile at 72 hpt showing the absolute value of the effects to identify parameters that are deviating from normality. Red line has a slope of 1 whereas the line blue line passes through the origin with a slope of the Lenth’s estimate of σ. (C) Comparison between actual and predicted Log(Vp) at 72 hpt with line of fit in red and the mean value in blue. (D) Residual of Log(Vp) vs predicted Log(Vp Vp at 72 hpt to check if the values are randomly scattered around zero (blue). (E) Residual vs row number visualization at 72 hpt with a line at 0 (blue). (F), (G), (H) and (I) are equivalent to (B), (C), (D), and (E) for viability. Surface plots for Log(Vp) and (J) pHelper and pRepCap. (K) pHelper and FectoVIR. (L) pRepCap, FectoVIR. (M) pHelper and pGOI. (N) pGOI and pRepCap. (O) pGOI and FectoVIR. Contour lines with Log(Vp) values at the bottom for all plots.

**Figure 3:**
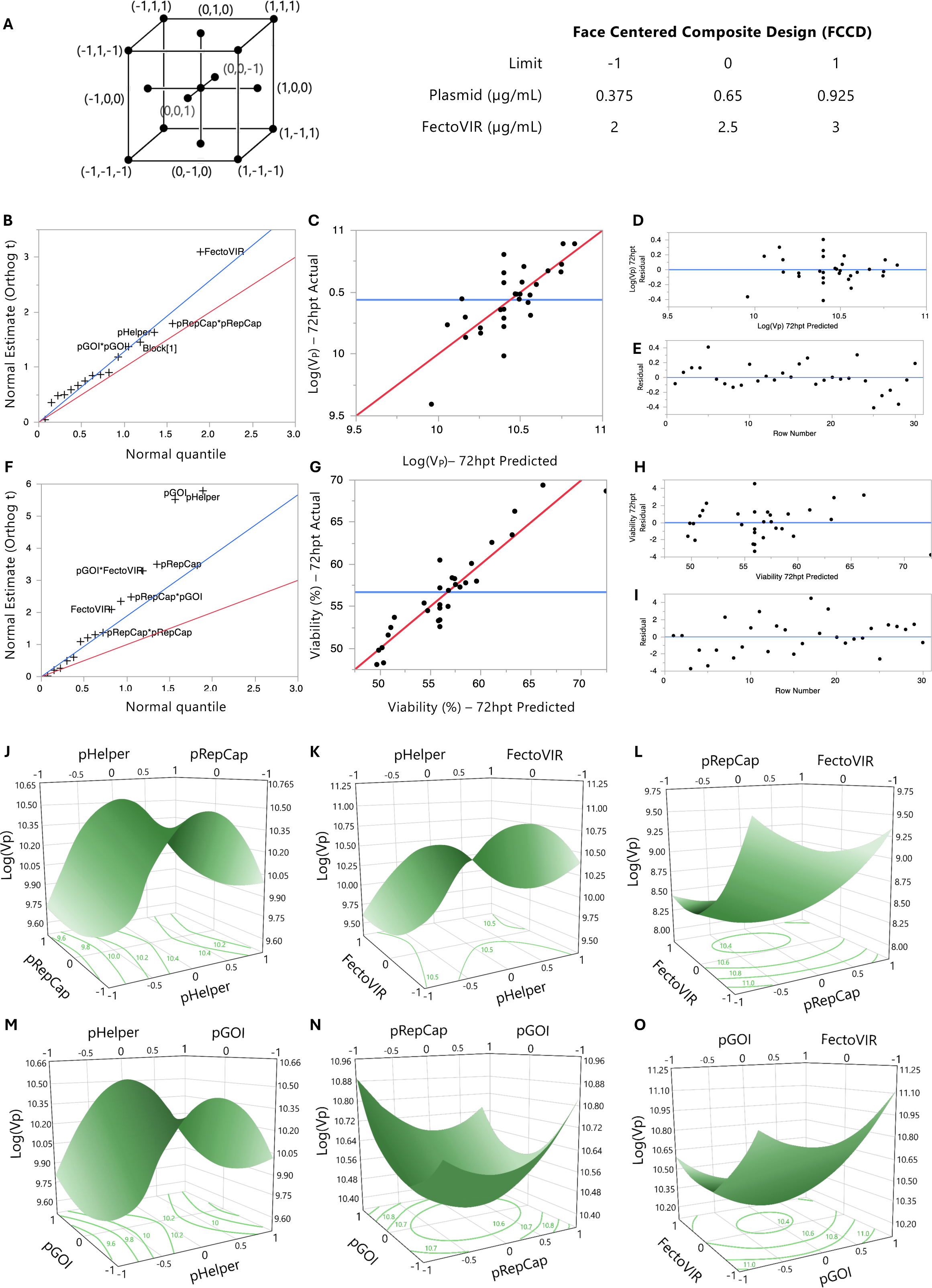
FCCD 72 hpt. (A) Graphic representation of experimental space of a 4-factors FCCD (left) and limits used for the model (right). (B) Normal estimate (Orthog t) against normal quantile at 72 hpt showing the absolute value of the effects to identify parameters that are deviating from normality. Red line has a slope of 1 whereas the line blue line passes through the origin with a slope of the Lenth’s estimate of σ. (C) Comparison between actual and predicted Log(Vp) at 72 hpt with line of fit in red and the mean value in blue. (D) Residual of Log(Vp) vs predicted Log(Vp) at 72 hpt to check if the values are randomly scattered around zero (blue). (E) Residual vs row number visualization at 72 hpt with a line at 0 (blue). (F), (G), (H) and (I) are equivalent to (B), (C), (D), and (E) for viability. Surface plots for Log(Vp) and (J) pHelper and pRepCap. (K) pHelper and FectoVIR. (L) pRepCap, FectoVIR. (M) pHelper and pGOI. (N) pGOI and pRepCap. (O) pGOI and FectoVIR. Contour lines with Log(Vp) values at the bottom for all plots.

For instance, for Log(Vp) at 72 hpt in the RCCD model (**Fig 2B**), Block1 and Block2 are significant and, therefore, this source of variability must be considered. Since the blocks were assigned based on the qPCR plate used, it was henceforth treated as a random effect. After defining blocking as a random effect, the model analysis using the restricted maximum likelihood method (REML) was repeated. We then compared the p-value, RMSE and R-square and the percentage of variability attributed to blocking for the three RSM models at both 48 and 72 hpt with and without blocking as a random effect (**Table S9**).

When analyzing viability as a response in each of our models, blocking was not required, as the samples were all quantified at the same time and did not suffer from the external source of uncontrolled noise coming from the qPCR plate. This was supported by two facts: a) the highest variability observed in viability due to blocking was 5.66% for FCCD at 72 hpt (**Table S4**), while in BBD and RCCD, the blocking only accounted for 0.87% and 0.00% of the variability, respectively (**Table S6, S2**) and b) the highest p-value for all models concerning viability was 0.0022 (**Table S9).**

Regarding the response of Log(Vp), blocking had a different impact on each model. For RCCD, blocking improved the model fitness increasing the R-square value from 0.57 to 0.72 at 48 hpt and from 0.29 to 0.81 for 72 hpt with a subsequent decrease in the p-value from 0.2526 to 0.1158 and 0.9396 to 0.246 respectively. Furthermore, the blocking accounted for 64.67% of the total variability at 72 hpt (**Table S2**).

Contrarily, in the FCCD, blocking worsened the model fitness at 48 hpt raising the p-value from 0.0597 to 0.2877 with a decrease in the R-square from 0.68 to 0.589. At 72 hpt, the obtained model fitness parameters were not relevantly different with an R-square value of 0.59 when blocking was applied versus 0.58. As expected, only 0.89% of the variability originated from the blocking at 72 hpt.

Finally, for the BBD, the introduction of blocking at 48 hpt was not largely different from the parameters obtained after blocking. For instance, the p-value and R-square with blocking were 0.0684 and 0.6516 whereas the same parameters without blocking were 0.0474 and 0.70. However, at 72 hpt, blocking was crucial to decreasing the p-value from 0.7885 to 0.4273 and to increase the R-square from 0.38 and 0.67757. The variability stemming from the blocking was 34.39% at 72 hpt.

The large disparities between the three models further affirmed the random nature of the uncontrolled factor coming from the different qPCR plates, underscoring the need for blocking in our experimental setup. Hence, blocking must not only be encouraged, but also considered a crucial factor when not all samples can be accommodated in a single qPCR run or any other equipment generating high intrinsic variability and uncontrolled noise.

We have demonstrated the need for blocking when uncontrolled factors are introduced, such as when using qPCR for rAAV quantification. It is crucial to emphasize that the failure to apply blocking when experimentally necessary could lead to misleading conclusions. This became evident when we individually assessed the effect of blocking on each RSM approach. Interestingly, FCCD appeared not to require blocking, as evidenced by the lack of relevant changes in the p-value with and without blocking, and the low variability introduced due to blocking (**Table S9**). Conversely, RCCD showed the opposite effect—the introduction of blocking increased the model fitness and was able to explain a high percentage of the data variability. Additionally, BBD only demanded blocking at 72 hpt. We hypothesize that the different impact of blocking on each RSM is attributable to its unpredictable random effect. Therefore, to prevent incorrect conclusions and suboptimal solutions, it is necessary to first statistically test if blocking is influencing the studied response when uncontrolled factors are involved. Only in the absence of such factors, blocking would not be recommended.

### 3.2. Different models work differently for each response

DOE approaches serve as invaluable tools in optimizing bioprocess conditions for multiple responses simultaneously. Our study highlights the variability in model efficiency among different DOE approaches when optimizing viability and volumetric productivity during rAAV production, as well as the variability in the identification of factors significantly affecting the studied responses.

Regarding the response of Log(Vp), we discovered significant variability in the ability of the models to identify significant factors. The only and most significant factor affecting response variability at 72 hpt in RCCD appears to be pHelper (p-value = 0.04) (**Table S2**), underscoring the importance of helper functions in rAAV productivity. In contrast, when using FCCD, the model could only identify the transfection reagent as significant (p-value = 0.01) (**Table S4**). BBD failed to identify any significant factors affecting Log(Vp) (**Table S6**), while employing a MD followed by a 2-factors FCCD provided insight into previously unstudied interactions, highlighting all three plasmids, the transfection reagent and total DNA amount as significant factors (**Table S8**).

Moreover, we observed variability in the model fitting for Log(Vp) among the studied approaches. MD followed by a 2-factors FCCD yielded the highest R-square (0.952) (**Table S8**) and was the only significant fitting (p-value = 0.0002) for this response. The R-square value for RCCD was the second highest, with a value of 0.82 (**Table S2**), followed by BBD and FCCD (0.68 and 0.59, respectively) (**Table S4 and S6**). In addition, all RSM models analyzed by REML exhibited high p-values (>0.25) (**Table S2, S4, S6**), compared to MD and 2-factors FCCD for optimizing rAAV volumetric productivity.

Similarly to Log(Vp), we observed variability between the studied models regarding the response of viability. In all cases, pGOI and 2-factors interactions between the transfection reagent and at least one of the plasmids, were identified as significant. pHelper and pRepCap were significant for viability in all models at 72 hpt, except BBD (**Table S2, S4, S6 and S8**). In the case of MD, all three plasmids were identified as significant (p-value<0.002) (**Table S8**), while in the coupled 2-factors FCCD, the transfection reagent, total DNA amount and the interactions between these two were all significant (p-value<0.01) (**Table S4**). However, lower variability was observed in the model fitting when studying viability, indicating that all models could be efficiently used for optimizing this response. In this case, all models presented significant p-values (<0.0003) (**Table S2,S4,S6 and S8**) and high R-square values. MD-2 factors FCCD had the highest R-square value (0.97) (T**able S8**), followed by BBD (R-square = 0.95) (**Table S6**), FCCD (R-square = 0.89) (**Table S4**) and last, RCCD (R square = 0.87) (**Table S2**).

Here we show that minor differences in the design of the experimental space can greatly affect the ability of the model to identify significant interactions between the factors and their effect in the studied responses. Interestingly, we observe that among RSM approaches, RCCD would be the most efficient option for optimizing rAAV productivity. However, it would be the least preferred option for viability optimization, as BBD showed the best performance among the 3 RSM approaches, highlighting that selecting and using a RSM method to analyze and optimize both responses at the same time can be challenging. Regarding only viability, BBD and MD followed by a 2-factors FCCD emerged as preferred options. However, for simultaneous optimization of both viability and Log(Vp), MD followed by a 2-factors FCCD outperformed the RSM approaches. These findings underscore the importance of selecting the appropriate DOE approach tailored to the specific response of interest in bioprocess optimization studies.

### 3.3. Significant factors for harvesting timepoint selection

The selection of an appropriate harvest time point is crucial, as it significantly impacts both the quantity and quality of the final product. Balancing our productivity metric with viability considerations is essential, as low viability levels may trigger protease release, potentially compromising product quality^38^. Our results indicate that different parameters should be considered when selecting the harvesting timepoint. For the 4-factor RCCD model, the highest Log(Vp) values are observed at 72 hpt, with high viability achievable at both 72 hpt and 48 hpt (**Fig 6**). When harvesting at 72 hpt, implementing a blocking method is crucial, as it significantly affects volumetric productivity (**Fig 2 B**). However, this trend is not observed at 48 hpt, where pGOI and pHelper play a more significant role in response variability (**Fig S1 B**). Plasmid interactions with the transfection reagent were identified as significant factors for viability at both 48 hpt and 72 hpt.

For the 4-factor FCCD model, the highest Log(Vp) and viability values were both achieved at 48 hpt (**Table S3**), indicating that 48 hpt would be the optimal harvest time point when using this model to analyze our system. The amount of the transfection reagent emerged as the most significant factor for volumetric productivity at both timepoints (**Table S3, S4).** The transfection reagent was significant only for viability at 48 hpt, with pGOI and pHelper affecting culture viability more than other factors (**Table S3, S4**). When optimizing Log(Vp) using BBD, the highest values were observed at 48 hpt, reaching 9.88 followed by high viability levels, above 75% (**Table S5**). At this timepoint, the transfection reagent appeared to be the most significant factor for both Log(Vp) and viability (**Fig S3 A, D**). At 72 hpt, similarly to RCCD, blocking was observed to be crucial for optimizing Log(Vp) (**Fig 2 B, Table S9**). However, viability was still observed mostly affected by the transfection reagent (**Fig 4 F**). Lastly, the highest Log(Vp) values, achieved with MD followed by a 2-factor FCCD were observed at 72 hpt, with viability levels exceeding 78% **(Fig 1, Table S8**), highlighting the efficacy of this model in finding optimal values for Log(Vp) while maintaining favorable viability levels. Notably, this model yielded the highest Log(Vp) values among all studied approaches, reaching 12.7 (**Fig 1 A – Fig 5**). Combining these approaches provided insight into new interactions and we were able to identify the transfection reagent, total amount of DNA, and their interactions as the most significant factors for both responses at 72 hpt (**Fig 5 A, E**). At 48 hpt, volumetric productivity and viability were mainly driven by the three plasmids involved in the triple transfection (**Fig S2 A, D**).

**Figure 4:**
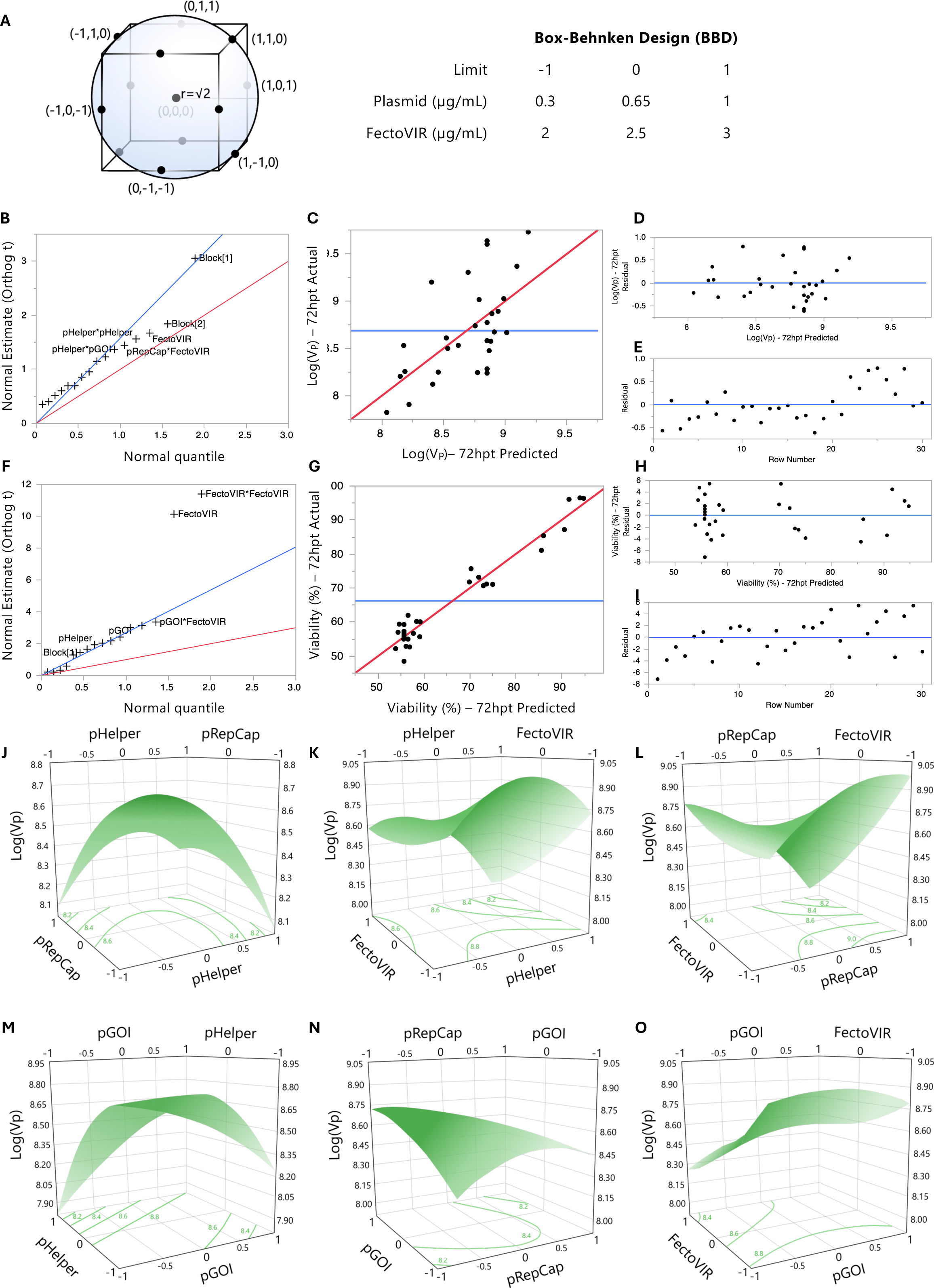
BBD 72 hpt. (A) Graphic representation of experimental space of a 4-factors BBD (left) and limits used for the model (right). (B) Normal estimate (Orthog t) against normal quantile at 72 hpt showing the absolute value of the effects to identify parameters that are deviating from normality. Red line has a slope of 1 whereas the line blue line passes through the origin with a slope of the Lenth’s estimate of σ. (C) Comparison between actual and predicted Log(Vp) at 72 hpt with line of fit in red and the mean value in blue. (D) Residual of Log(Vp) vs predicted Log(Vp) at 72 hpt to check if the values are randomly scattered around zero (blue). (E) Residual vs row number visualization at 72 hpt with a line at 0 (blue). (F), (G), (H) and (I) are equivalent to (B), (C), (D), and (E) for viability. Surface plots for Log(Vp) and (J) pHelper and pRepCap. (K) pHelper and FectoVIR. (L) pRepCap, FectoVIR. (M) pHelper and pGOI. (N) pGOI and pRepCap. (O) pGOI and FectoVIR. Contour lines with Log(Vp) values at the bottom for all plots.

**Figure 5:**
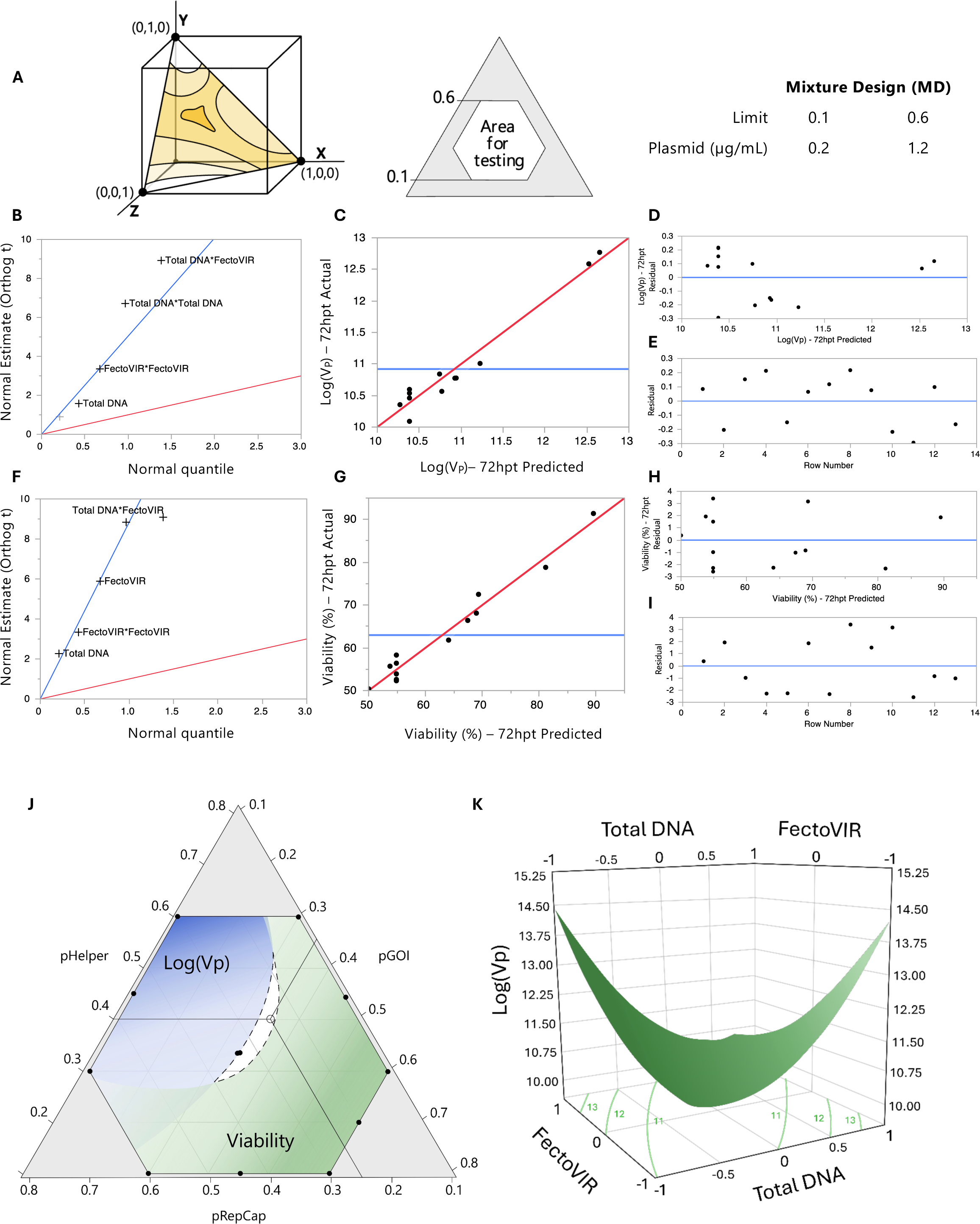
MD 72 hpt. (A) Graphic representation of experimental space of a 3-factors MD (left) and limits used for the model (right). (B) Normal estimate (Orthog t) against normal quantile at 72 hpt showing the absolute value of the effects to identify parameters that are deviating from normality. Red line has a slope of 1 whereas the line blue line passes through the origin with a slope of the Lenth’s estimate of σ. (C) Comparison between actual and predicted Log(Vp) at 72 hpt with line of fit in red and the mean value in blue. (D) Residual of Log(Vp) vs predicted Log(Vp) at 72 hpt to check if the values are randomly scattered around zero (blue). (E) Residual vs row number visualization at 72 hpt with a line at 0 (blue). (F), (G), (H) and (I) are equivalent to (B), (C), (D), and (E) for viability. (J) Experimental space showing both how Log(Vp) in blue and viability in green change at different ratios. Threshold for Log(Vp) set at 10.4 and for viability at 53%. (K) Surface plot for Log(Vp), total DNA and FectoVIR. Contour lines with Log(Vp) values at the bottom.

Our data revealed that each DOE model offered a unique perspective on the optimal harvest time point for rAAV production. These findings underscore the complexity of harvest time point selection in rAAV production, emphasizing the need to properly select the design approach to avoid misleading conclusions regarding both productivity and viability metrics.

### 3.4. Different optimal ratios were obtained using each model

We derived the optimal plasmid ratio and FV concentration from each strategy using the generated models to maximize Log(Vp) at 72 hpt. For some models, visualizing the optimal regions of the experimental space was clear (**Fig 2 K, Fig 2 L, Fig S3 I-K**).

However, when considering both responses at the same time and for some models like FCCD and BBD, the presence of saddle points (**Fig 3 J, K, L**) required the use of the desirability function (**Eq 4, 5**) to determine the optimal ratio for maximum productivity. We prioritized Log(Vp) with a desirability of 1 and viability with a desirability of 0.3.

Ultimately, the suggested optimal concentration of each plasmid and FV derived from each model was different. For instance, the optimal DNA concentration ratio for the 4-factors RCCD was 0.72:0.93:0.38 (pHelper:pRepCap:pGOI) with a FV concentration of 2 µg/mL, whereas for the 4-factors FCCD was 0.56:0.38:0.38 with a total of 2 µg/mL of FV. For the BBD, the optimal ratio was 0.66:0.64:0.38 coupled with 2.5 µg/mL of FV. The only value shared by the three RSM models was the 0.38 µg/mL of pGOI concentration. However, whereas FCCD required the same concentration of pRepCap, both BBD and RCCD demanded a higher concentration of the remaining plasmids, with pRepCap being the one with the highest concentration in RCCD (0.99 µg/mL) and pHelper being the highest concentration in BBD (0.66 µg/mL). Moreover, the optimal total amount of DNA in each model ranged from 2.09 µg/mL in RCCD to 1.31 µg/mL for FCCD, with BBD falling in the middle at 1.67 µg/mL. The calculated ratio between DNA and FV was 1.19 for RCCD, and despite the different plasmid ratios, BBD and FCCD had similar DNA:FV ratios of 0.67 and 0.66, respectively. Regarding the mixture design, the optimal concentration ratio was 0.80:0.70:0.50 with a total amount of DNA of 2 µg/mL and 2 µg/mL of FV. From the optimal plasmid ratio, the subsequent FCCD found the optimal ratio DNA:FV at 2.37.

All optimal values were successfully validated **(Fig 6)**, showing that the selection of suboptimal DOE models can provide misleading optimal values. For instance, the optimal ratio and consequently highest Log(Vp) achieved with BBD was on average 10.4 whereas the maximum Log(Vp) reached in MD followed by FCCD was 12.77 (**Fig 1, 6**) proving how the optimal value obtained from BBD was indeed suboptimal.

**Figure 6:**
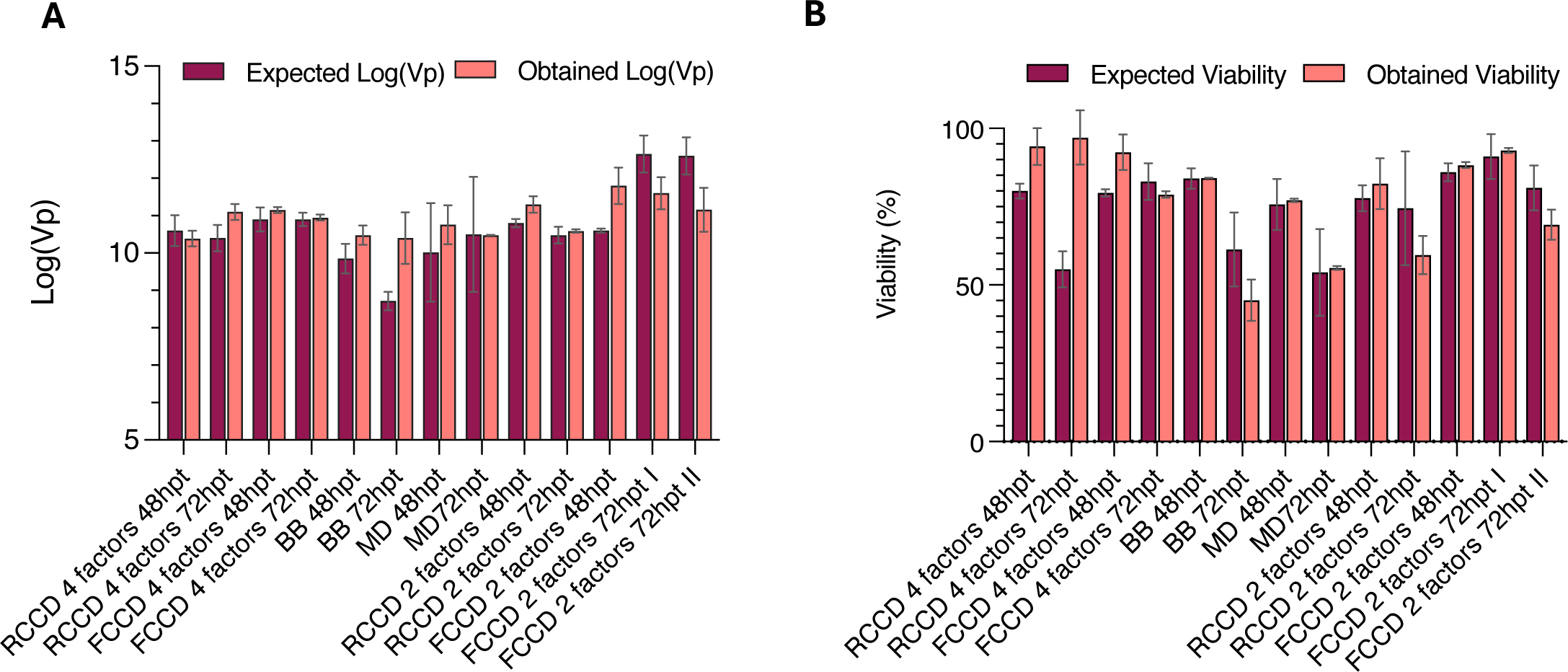
Validations. Predicted and actual experimental values for all validated conditions at 48 and 72 hpt for all models regarding (A) Log(Vp) and (B) viability.

### 3.5. Coupling MD with FCCD outperformed RSM approaches

Among the various DOE approaches studied, MD followed by a 2-factors FCCD emerged as the most efficient in enhancing both viral productivity and viability. This approach yielded the highest Log(Vp) values and required only 25 experimental runs in total, compared to the 4-factors RSM models that consisted of 30 runs. This indicates high efficiency in experimental resource utilization **(Fig 5 A**) and reduced variability due to the absence of the need for blocking. Using a D-optimal MD, we determined that the optimal plasmid ratio was 0.4:0.35:0.25 for pHelper:pGOI:pRepCap, respectively. After fixing this optimal ratio, explored whether a 2-factors RCCD or FCCD could further optimize the total DNA amount and the transfection reagent volume. The FCCD emerged as the preferred option, demonstrating better model fitting (**Fig S5**) and complementing the mixture design to identify transfection conditions that further improved Log(Vp) and viability values to 12.7 and 78.8%, respectively (**Fig 6**). Moreover, the 2-factors FCCD model after MD exhibited satisfactory R-square and adjusted R-square model fitting values at 72 hpt (**Fig 5**). Importantly, this was the only approach to yield significant p-values for both responses (0.0002 for Log(Vp) and <.0001 for viability), unlike the RSM designs. Validation runs further confirmed the model’s fitting and accuracy in predicting viability and Log(Vp) values at 48 and 72 hpt (**Fig 6**).

The 2-factor FCCD after MD allowed for the specific study of the effects of total DNA in the system. Here, we observed two areas for optimal Log(Vp) at 72 hpt. The first area corresponded to the lowest concentration of DNA (0.93 µg/mL) and the highest concentration of FV (2.7 µg/mL), while the second area occurred with the highest DNA concentration (3.06 µg/mL) and the lowest FV concentration (1.29 µg/mL). To understand the appearance of these opposing solutions, we investigated the dynamics of the transfection system.

There are two different strategies to increase the number of plasmid copies in the cell: increase the plasmid concentration in the culture or increase the transfection reagent. However, high concentrations of transfection reagent can be cytotoxic and trigger cellular death^41^. Therefore, it is crucial to find the ideal concentrations to maximize endogenous protein synthesis pathways while not compromising viability.

For instance, when DNA amount and FV are at their highest, it could potentially result in the highest amount of DNA in the cell, the highest cytotoxic protein synthesis and FV toxicity.

Therefore viability and Log(Vp) drastically decreased (**Fig 5, Table S8**). This is supported by the two opposing areas for optimal Log(Vp) in the 2-factors FCCD. Additionally, the cytotoxic effect of FV can be observed, as increasing concentrations lead to a steady decrease in viability. The obtained optimal DNA concentration of 1 µg of DNA per mL of culture agrees with the manufacturer recommendations for optimal expression. However, the observed ratios to maximize Log(Vp) were 3.06:1.29 and 0.93:2.70 DNA:FV, diverging from the generally recommended 1:1. Using a MD-FCCD approach allows the analysis of the DNA-transfection reagent interaction showing that when DNA concentration is close to 1 µg/mL, a high concentration of FV will be necessary to ensure high productivity. However, when DNA concentration is high, a FV concentration close to 1.29 µg/mL should suffice to achieve maximum productivity.

Alternatively, in the 2-factors RCCD after the MD, FV being toxic at high concentrations could also be observed (**Table S8, S10**). However, on the opposite end, with a low concentration of FV, Log(Vp) did not suffer any relevant change, regardless of the quantity of DNA. This response proved that FCCD is more suitable than to better study all biological phenomena and provide better-defined ratios.

The primary objective of this work was to provide necessary tools to address challenges encountered in optimizing current rAAV manufacturing processes. When selecting an appropriate model, a thorough understanding of the biological context is crucial. In the triple transfection process, there are many factors that affect the final outcome, and they should be appropriately analyzed to avoid masking and confounding effects in the models. The cell density is crucial for transfection-based processes. However, the biological reason behind the effects of the cell density in the process is substantially different to the biological reason to define a certain plasmid ratio, for instance. The cell density effect (CDE) acts as the limiting factor for cell density, imposing a threshold beyond which transfection efficiency and cell-specific productivity decrease. This is a common phenomenon shared across different cell line platforms and expression methods, unrelated to the expression of AAV elements^4,39^. Therefore, to thoroughly study the interactions between AAV elements and reduce variables to optimize the experimental design, the cell density can be fixed at a value that is not yet affected by the CDE in our system.

Subsequently, the three plasmids and the transfection reagent can be defined as independent factors and were treated as such in all RSM designs (RCCD, FCCD, and BBD), resulting in two different ratios: pGOI:pHelper:pRepCap and DNA:Transfection Reagent. However, the total DNA amount added to the cell can lead to independent effects, solely driven by the DNA total concentration. These effects are overlooked when using RSM approaches. To mitigate this, the total DNA concentration can be fixed as one of the variables to study. Thus, a relationship between all plasmid concentrations will emerge in the form of a ratio, and they can no longer be treated as independent variables, as an increase in one would cause a proportional decrease in the others. This could lead to suboptimal solutions or misleading estimate factors when analyzing these models. Consequently, the MD is necessary to successfully study the system. It separates the analysis of the plasmid interactions and their effects on the studied responses, treating the three plasmids as parts of a total mixture. The biological effects of expressing the different AAV elements and their interactions with cellular homeostasis are separate from the effects of just a high transfected DNA amount, regardless of the coded proteins. The first step of the MD studies the expressed proteins and the significance of their effects on the studied responses and the process. This is the main biological reasoning behind the optimal plasmid ratio. Subsequently, this optimal ratio is used to study the separate effects of the total amount of DNA and transfection reagent concentration. Although still related to the coded proteins, these effects have a separate biological effect, shared by any transfection-based process. Using 4-factors RSM approaches, these two biological scenarios are mixed, leading to confounded effects. The coupling of MD-RSM enabled us to better analyze biological phenomena by separating variables according to their biological relevance. This approach provides insight into the role of the total DNA amount, something that is impossible when using RSM methodologies, as described above.

One of the most crucial variables in the process is the GOI, whose effects and interactions in the cell will substantially vary for each rAAV-based product. Unless engineered not to be expressed in the host cell line^40^, the GOI will have an impact on the cell metabolism and physiology, with potential interactions with the rest of the AAV elements that the DOE model should be able to identify and analyze. Moreover, the limits of the experimental space to study new GOIs will be constrained to their effect on the cell. Since the behavior of the GOI may differ depending on the final product, we anticipate that testing other GOIs will yield different responses. Our results showed that a D-optimal quadratic MD approach is the most suitable design to study these effects, as it is able to focus on plasmid interactions better than RSM approaches.

**Supplementary Figure S1:**
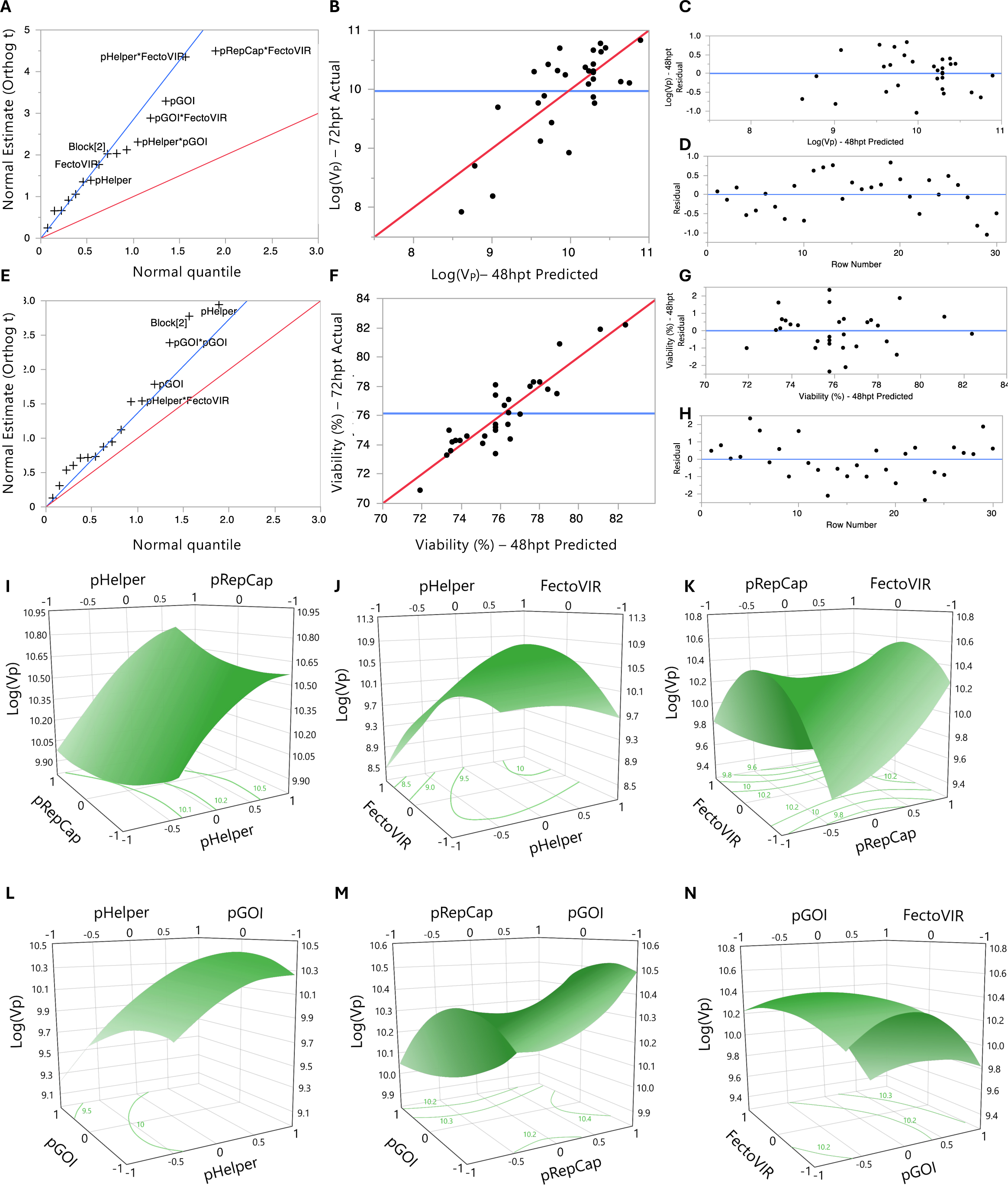
RCCD 48 hpt. (A) Normal estimate (Orthog t) against normal quantile at 48 hpt showing the absolute value of the effects to identify parameters that are deviating from normality. Red line has a slope of 1 whereas the line blue line passes through the origin with a slope of the Lenth’s estimate of σ. (B) Comparison between actual and predicted Log(Vp) at 48 hpt with line of fit in red and the mean value in blue. (C) Residual of Log(Vp) vs predicted Log(Vp) at 48 hpt to check if the values are randomly scattered around zero (blue). (D) Residual vs row number visualization at 48 hpt with a line at 0 (blue). (E), (F), (G) and (H) are equivalent to (A), (B), (C), and (D) for viability. Surface plots for Log(Vp) and (I) pHelper and pRepCap. (J) pHelper and FectoVIR. (K) pRepCap, FectoVIR. (L) pHelper and pGOI. (M) pGOI and pRepCap. (N) pGOI and FectoVIR. Contour lines with Log(Vp) values at the bottom for all plots.

**Supplementary Figure S2:**
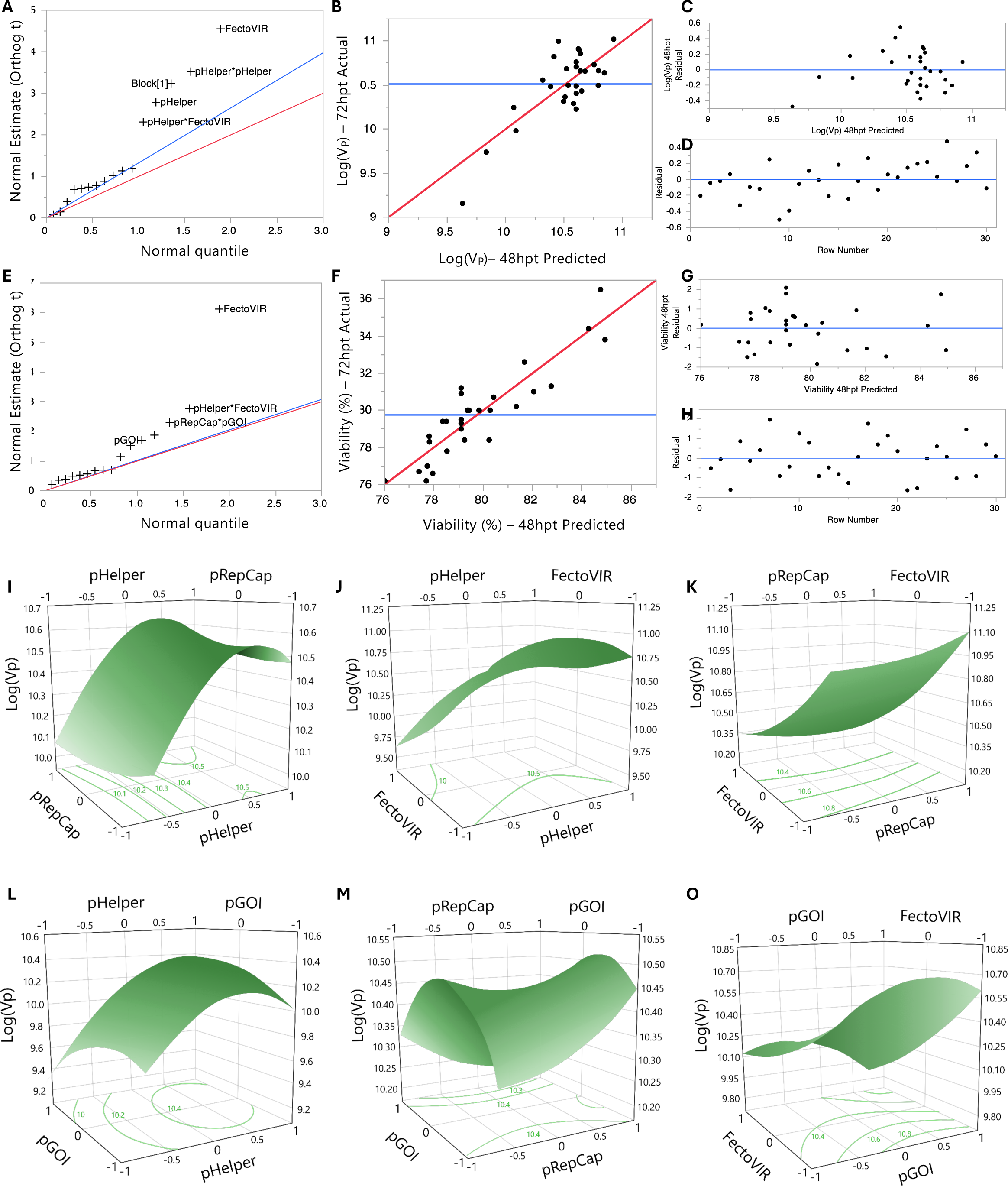
FCCD 48 hpt. (A) Normal estimate (Orthog t) against normal quantile at 48 hpt showing the absolute value of the effects to identify parameters that are deviating from normality. Red line has a slope of 1 whereas the line blue line passes through the origin with a slope of the Lenth’s estimate of σ. (B) Comparison between actual and predicted Log(Vp) at 48 hpt with line of fit in red and the mean value in blue. (C) Residual of Log(Vp) vs predicted Log(Vp) at 48 hpt to check if the values are randomly scattered around zero (blue). (D) Residual vs row number visualization at 48 hpt with a line at 0 (blue). (E), (F), (G) and (H) are equivalent to (A), (B), (C), and (D) for viability. Surface plots for Log(Vp) and (I) pHelper and pRepCap. (J) pHelper and FectoVIR. (K) pRepCap, FectoVIR. (L) pHelper and pGOI. (M) pGOI and pRepCap. (N) pGOI and FectoVIR. Contour lines with Log(Vp) values at the bottom for all plots.

**Supplementary Figure S3:**
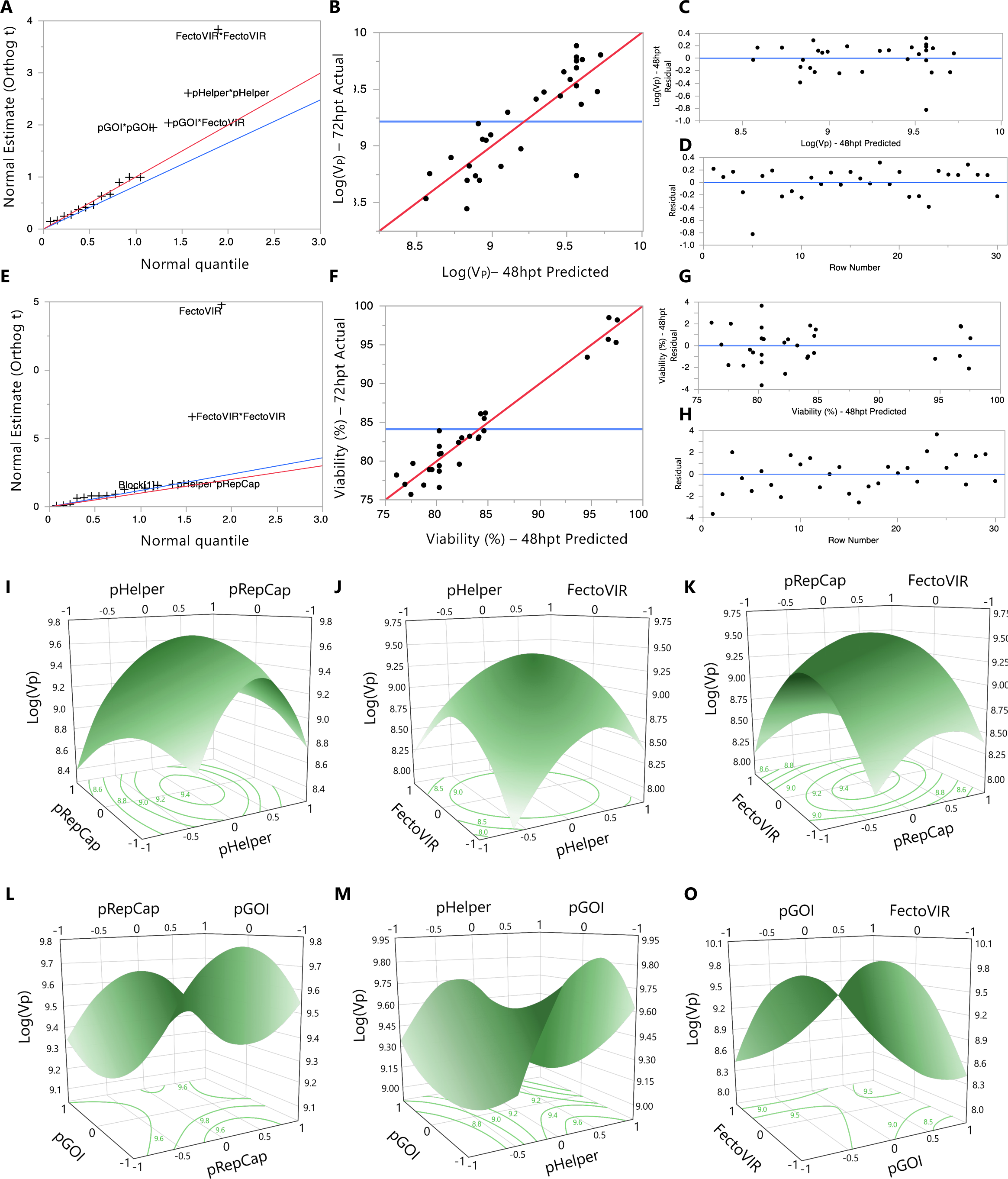
BBD 48 hpt. (A) Normal estimate (Orthog t) against normal quantile at 48 hpt showing the absolute value of the effects to identify parameters that are deviating from normality. Red line has a slope of 1 whereas the line blue line passes through the origin with a slope of the Lenth’s estimate of σ. (B) Comparison between actual and predicted Log(Vp) at 48 hpt with line of fit in red and the mean value in blue. (C) Residual of Log(Vp) vs predicted Log(Vp) at 48 hpt to check if the values are randomly scattered around zero (blue). (D) Residual vs row number visualization at 48 hpt with a line at 0 (blue). (E), (F), (G) and (H) are equivalent to (A), (B), (C), and (D) for viability. Surface plots for Log(V_P_) and (I) pHelper and pRepCap. (J) pHelper and FectoVIR. (K) pRepCap, FectoVIR. (L) pHelper and pGOI. (M) pGOI and pRepCap. (N) pGOI and FectoVIR. Contour lines with Log(Vp) values at the bottom for all plots.

**Supplementary Figure S4:**
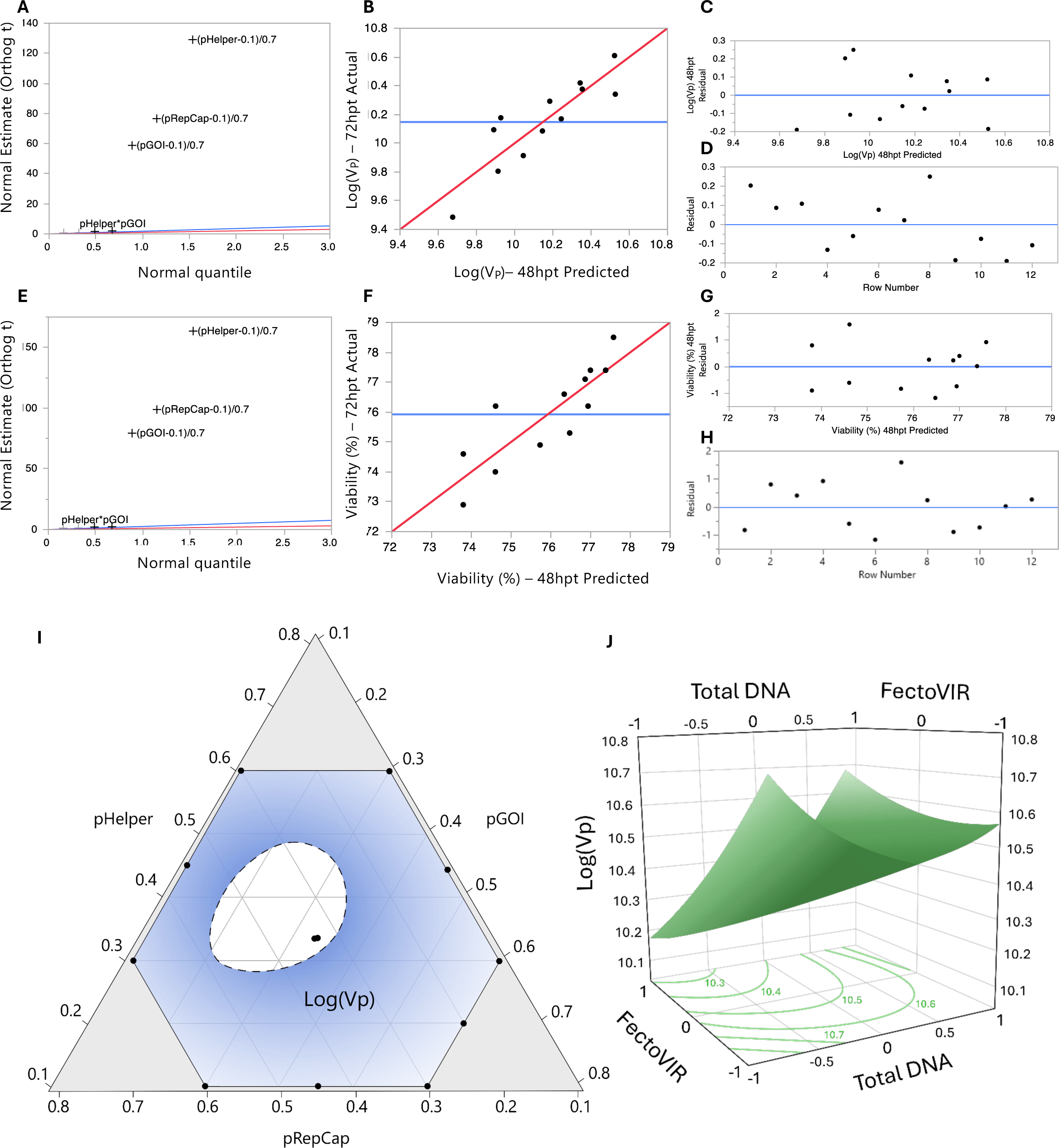
MD 48hpt. (A) Normal estimate (Orthog t) against normal quantile at 48 hpt showing the absolute value of the effects to identify parameters that are deviating from normality. Red line has a slope of 1 whereas the line blue line passes through the origin with a slope of the Lenth’s estimate of σ. (B) Comparison between actual and predicted Log(V_P_) at 48 hpt with line of fit in red and the mean value in blue. (C) Residual of Log(V_P_) vs predicted Log(V_P_) at 48 hpt to check if the values are randomly scattered around zero (blue). (D) Residual vs row number visualization at 48 hpt with a line at 0 (blue). (E), (F), (G) and (H) are equivalent to (A), (B), (C), and (D) for viability. (I) Experimental space showing how Log(Vp) in blue change at different ratios. Threshold for Log(Vp) set at 10.4. (J) Surface plot for Log(Vp), total DNA and FectoVIR. Contour lines with Log(Vp) values at the bottom.

**Supplementary Figure S5:**
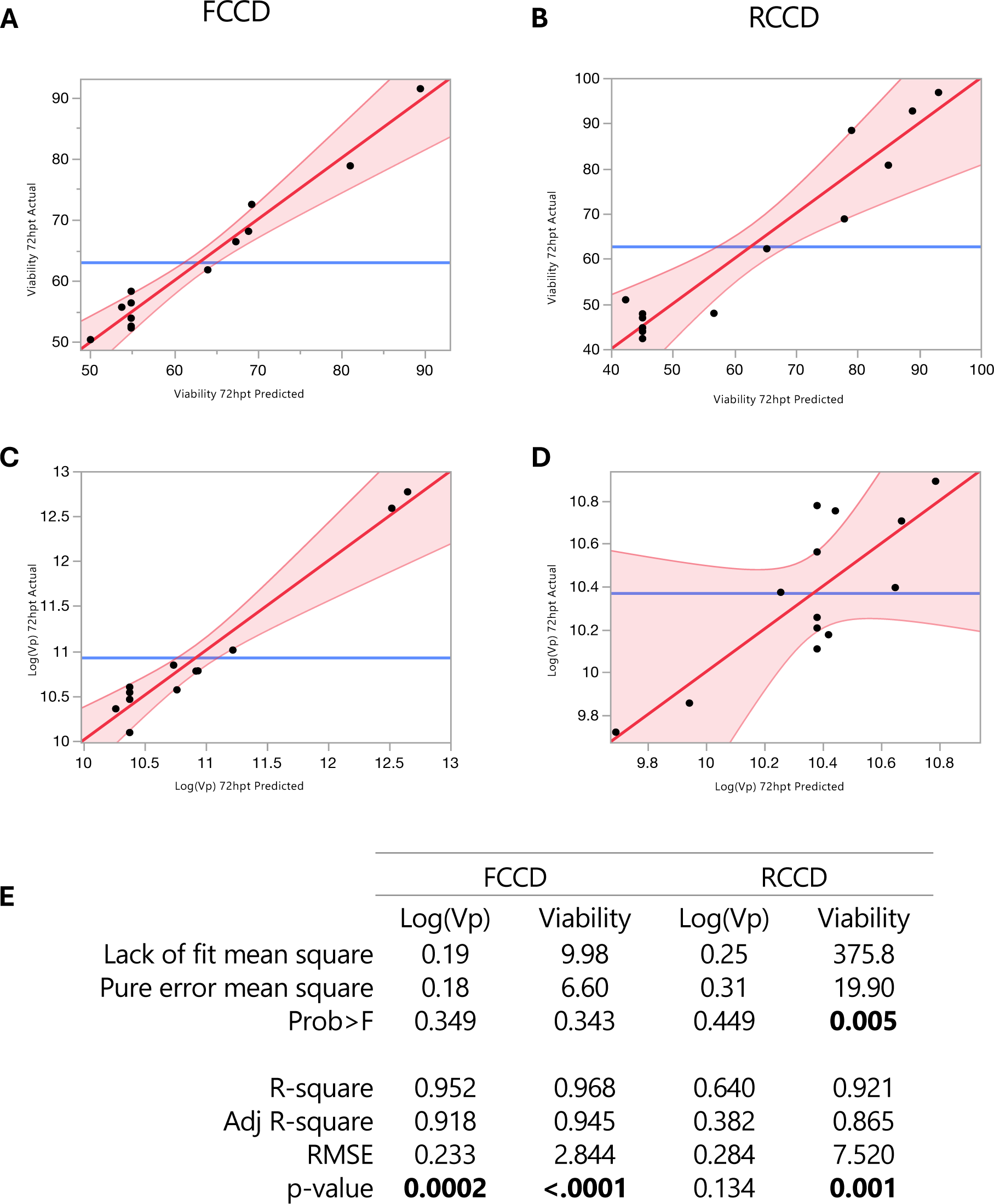
Comparison 2-factors FCCD vs RCCD. (A) Actual by Predicted plot for Viability at 72 hpt, FCCD. (B) Actual by Predicted plot for Viability at 72 hpt, RCCD. (C) Actual by Predicted plot for Log(Vp) at 72 hpt, FCCD. (D) Actual by Predicted plot for Log(Vp) at 72 hpt, RCCD. (E) Lack of fit and Summary of fit analyses for FCCD and RCCD.

**Supplementary Table S1: Statistical parameters RCCD 48 hpt.** Matrix with coded limits for each run with the corresponding obtained response values for Log(Vp) and viability. Statistical parameters from REML analysis to assess model fitness and estimate significance are shown.

**Supplementary Table S2: Statistical parameters RCCD 72 hpt.** Matrix with coded limits for each run with the corresponding obtained response values for Log(Vp) and viability. Statistical parameters from REML analysis to assess model fitness and estimate significance are shown.

**Supplementary Table S3: Statistical parameters FCCD 48 hpt.** Matrix with coded limits for each run with the corresponding obtained response values for Log(Vp) and viability. Statistical parameters from REML analysis to assess model fitness and estimate significance are shown.

**Supplementary Table S4: Statistical parameters FCCD 72 hpt.** Matrix with coded limits for each run with the corresponding obtained response values for Log(Vp) and viability. Statistical parameters from REML analysis to assess model fitness and estimate significance are shown.

**Supplementary Table S5: Statistical parameters BBD 48 hpt.** Matrix with coded limits for each run with the corresponding obtained response values for Log(Vp) and viability. Statistical parameters from REML analysis to assess model fitness and estimate significance are shown.

**Supplementary Table S6: Statistical parameters BBD 72 hpt.** Matrix with coded limits for each run with the corresponding obtained response values for Log(Vp) and viability. Statistical parameters from REML analysis to assess model fitness and estimate significance are shown.

**Supplementary Table S7: Statistical parameters MD 48 hpt.** Matrix with coded limits for each run with the corresponding obtained response values for Log(Vp) and viability. Statistical parameters from REML analysis to assess model fitness and estimate significance are shown.

**Supplementary Table S8: Statistical parameters MD 72 hpt.** Matrix with coded limits for each run with the corresponding obtained response values for Log(Vp) and viability. Statistical parameters from REML analysis to assess model fitness and estimate significance are shown.

**Supplementary Table S9: Blocking versus non-blocking.** Fitness parameters for each model at 48 and 72 hpt RSM models with and without blocking.

**Supplementary Table S10: Comparison between 2-factors RCCD and FCCD after MD.** Matrix with coded limits for each run and corresponding obtained response values for Log(Vp) and viability. Statistical parameters from REML analysis to assess model fitness and estimate significance are shown.

## CONFLICT OF INTEREST

All authors declare no conflict of interest.

## AUTHOR CONTRIBUTIONS

KT and DCT: Investigation, visualization, writing original draft, review and editing. LKN: Supervision and review. JLG: Conceptualization, visualization, supervision, writing original draft, review and editing.

## ACKNOWLEDGEMENTS AND FUNDING

The authors would like to acknowledge generous support by Novo Nordisk Foundation. This work was supported by NNF20CC0035580 and NNF20SA0066621. LKN is supported by NNF14OC0009473. JLG is supported by NNF22OC0078741 and Marie Skłodowska-Curie Actions (MSCA) Postdoctoral Fellowship 101105465. KT is supported by a DTUQ Alliance PhD grant.

## Notes

### Competing Interest Statement

The authors have declared no competing interest.

